# Prefoldin facilitates the action of FACT on macroH2A1-containing chromatin during transcription elongation

**DOI:** 10.64898/2026.11.15.766783

**Authors:** Sara Fontalva, Elena Sánchez-Escabias, Esperanza Granado, Xenia Peñate, Rocío Cañete, Álvaro Gallego-Martínez, Vicent Pelechano, María C. Muñoz-Centeno, José C. Reyes, Sebastián Chávez

## Abstract

Prefoldin is one of the chaperone complexes that participate in protein homeostasis. Although it was initially described as a co-translational cochaperon that captures newly synthesized polypeptides and facilitates their folding by transferring them to class II chaperonins, nuclear functions have also been reported, including its contribution to transcription elongation and chromatin dynamics from budding yeast to mammalian cells. In this work, we have performed a proteomic search for prefoldin interactors in chromatin. Among others, we detected its physical association to FACT, the main histone chaperon that operates during transcription elongation. Lack of prefoldin provokes phenotypes associated to FACT perturbation, like chromatin-integrity defects in transcribed gene bodies, and derepression of cryptic transcription. It also leads to the specific alteration of FACT transcriptional dynamics in prefoldin-regulated genes. Lack of prefoldin also provokes co-transcriptional defects in histone dynamics, particularly in the eviction of macroH2A1, being this non-canonical histone a prefoldin-interactor itself. Finally, we show that these physical and functional connections with FACT and macroH2A1 respond to a single molecular phenomenon, and propose a model in which prefoldin would play a bridging role, enabling FACT to drive co-transcriptional eviction of macroH2A1-containing nucleosomes.

## Introduction

Prefoldin is a highly conserved hexameric co-chaperone present in archaea and eukaryotes. In eukaryotic cells, the canonical complex is composed of six distinct subunits (PFDN1-6), arranged into two α-type subunits (PFDN3 and PFDN5) and four β-type subunits (PFDN1, PFDN2, PFDN4, and PFDN6), forming a characteristic jellyfish-like structure [1].

Prefoldin is best known for its cytoplasmic role in protein homeostasis, where it captures unfolded or newly synthesized polypeptides and facilitates their folding by transferring them to the CCT/TRiC complex, an ATP-dependent class II chaperonin [2,3]. However, prefoldin is also present in the nucleus [4], where individual human prefoldin subunits have been implicated in functions that are fundamentally different from its cytoplasmic role [5–8]. In Saccharomyces cerevisiae, the canonical prefoldin complex was shown to associate with the coding regions of actively transcribed genes in a transcription-dependent manner and to contribute to RNA polymerase II elongation [9]. Importantly, prefoldin also contributes to co-transcriptional chromatin dynamics, including histone eviction during transcription elongation [9]. In the case of human cells, it has been demonstrated that human prefoldin associates with active chromatin and displays a distribution pattern similar to RNA polymerase II, locally influencing gene expression and co-transcriptional pre-mRNA splicing [10]. In response to serum starvation, prefoldin is required for the optimal regulation of a wide number of genes [10]. Together, these findings broaden the functional landscape of prefoldin, which appears as a regulator of transcription-associated chromatin dynamics.

In this work, we addressed a proteomic search for prefoldin interactors in the chromatin environment. Among them, we found the FACT complex, the main histone chaperone acting during transcription elongation to preserve chromatin integrity [10], and macroH2A1, a non-canonical histone involved in transcriptional repression [12]. We also found that lack of prefoldin provokes alterations in FACT and macroH2A1 co-transcriptional dynamics, and obtained evidence of prefoldin to play a bridging role in connecting FACT and macroH2A1 eviction during transcription.

## Results

### The canonical prefoldin complex associates to elongating RNApol II in the chromatin environment

The first objective of our work was to explore the set of physical interactors of prefoldin in the transcriptionally active chromatin environment. In order to address this specific subpopulation of cellular prefoldin, we optimized a protocol to purify prefoldin-containing protein complexes from chromatin (Fig 1A). After partially solubilizing the chromatin fraction of cells expressing a PFDN5-GFP fusion with benzonase, we ultracentrifuged it to remove the highly dense chromatin and used the supernatant to perform affinity purification through a GFP-Trap matrix. Figure 1B shows that the final pull-down fraction was highly enriched in PFDN5-GFP, free of cytoplasmic markers like tubulin, and without excess levels of histones, indicating that the pull-down fraction was depleted of unspecific contaminating chromatin.

**Figure 1:**
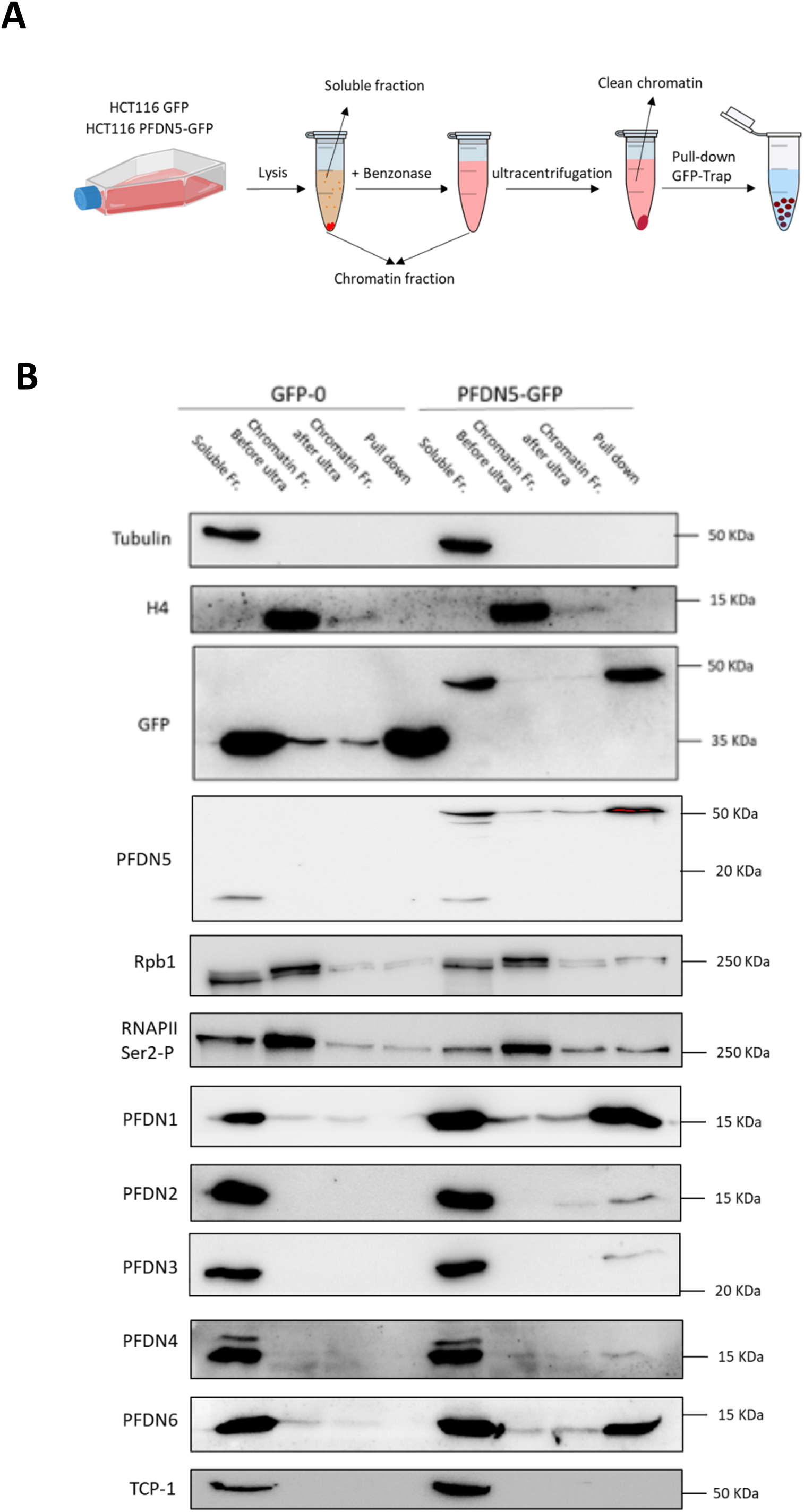
All canonical prefoldin complex subunits are present in chromatin and associates to elongating RNApol II. A) Schematic representation of the experimental procedure of the GFP pull-down experiments performed to isolate prefoldin-interacting proteins from the chromatin fraction. Human HCT116 cells with exogenous GFP or with the exogenous PFDN5-tagged GFP construct were grown. The cells pellets were lysed and centrifuged to separate two fractions: the supernatant, mainly corresponding to the cytoplasm, and the chromatin fraction in the pellet. The latter was digested for 3 hours with benzonase, and clarified by ultracentrifugation. Following this, the chromatin fraction was ready to perform affinity purification using GFP-Trap. **B** Western blots of the PFDN5-GFP pull-down experiments and the corresponding GFP negative controls. The membranes were incubated with antibodies against the indicated proteins.

We did find RNApol II in the pull-down fraction, particularly the slower IIO form of POLR2A, in comparison to a non-fused GFP control. Accordingly, we also detected enrichment in Ser2-phosphorylated POLR2A, a characteristic mark of the elongating form of RNApol II (Fig 1B).

In addition, we also detected the other five subunits of the canonical prefoldin complex in the PFDN5-GFP pull-down fraction. In contrast, no presence of these subunits was found in the unfused GFP control. We concluded that we had obtained optimal material to identify potential interactors of prefoldin in the context of active transcription.

### Chromatin prefoldin physically interacts with the FACT complex

Six independent pull-down fractions of PFDN5-GFP, together with six unfused GFP controls, were analysed by MS/MS tandem mass spectrometry. The comparison of PFDN5-GFP pulldowns and GFP controls showed significant enrichment for the six prefoldin subunits, which clustered together, confirming the presence of its canonical complex in human chromatin (Fig 2A). No other polypeptide was present with such robustness in the PFDN5-GFP fractions, that exhibit a rather high heterogeneity. However, several proteins did undergo significant enrichment in the six preparations, considered as a whole, in comparison to the unfused GFP controls (Fig 2A). Gene ontology analysis of their molecular functions showed that the most significantly enriched categories were related to RNA-binding proteins and enzymes acting on RNA (Table S1). Nucleosome-binding proteins were also significantly enriched (Table S1). Among them, we found SSRP1 and SUPT16H, the two subunits of the FACT histone chaperon, which also clustered together (Fig 2A).

**Figure 2:**
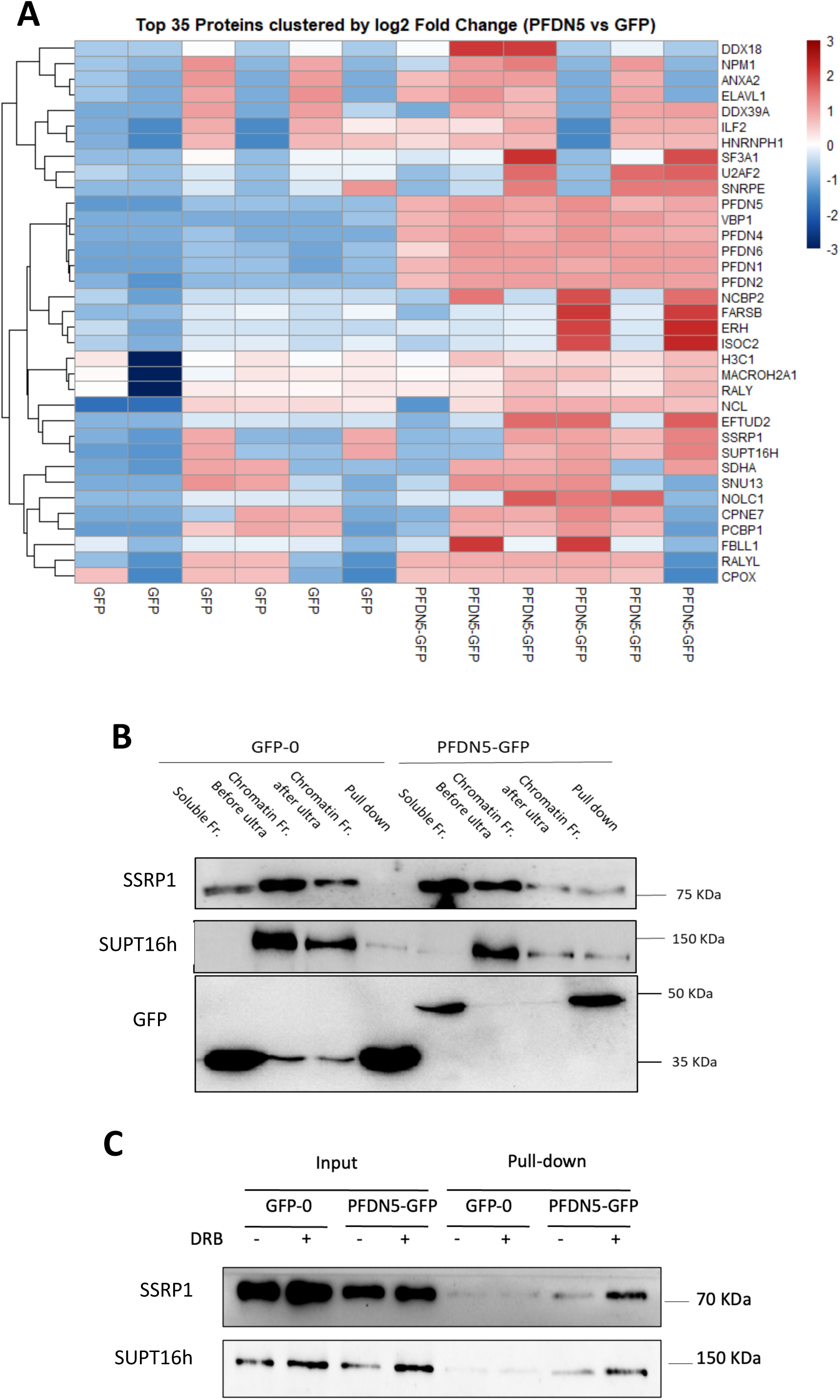
Chromatin prefoldin associates to the FACT complex in the context of transcription. A) Label-free quantitation of PFDN5-GFP and GFP pull-down mass spectrometry analyses. Total intensity data obtained with MSFragger were clustered by log2 PFDN5-GFP vs GFP fold change. B) The two subunits of the FACT complex are enriched in PFDN5-GFP pull-down experiments. Western blots of SSRP1 and SUPT16h (see Fig 1 for details). C) The interaction of PFDN5 with the FACT complex takes place before release from RNApol II promoter-proximal pausing. Western blot experiment of SSRP1 and SUPT16h after PFDN5-GFP pulldowns under DRB 100 µM treatment (+) or without DRB (-).

We confirmed the progressive enrichment of these two polypeptides across the prefoldin purification procedure described above (Fig 2B), without significant expression differences in the PFDN5-GFP context (Fig S1A). We also detected prefoldin associated to FACT when performing pulldown experiments in the opposite direction (Fig S1B). Finally, we found that prefoldin-FACT interaction takes place early, as the levels of FACT associated to PFDN5-GFP increased in the presence of the transcription-inhibitory drug DRB (Fig 2C). This result indicates that prefoldin-FACT interaction seems to be already occurring during early, before the release of RNApol II from promoter-proximal pausing.

### Lack of prefoldin alters FACT transcriptional dynamics and impairs chromatin integrity of transcribed genes

As we mentioned in the Introduction, prefoldin depletion provokes severe alterations in the transcriptomic response to serum starvation [10]. We decided to test whether lack of prefoldin also altered FACT dynamics in the same conditions. Since FACT recruitment to transcribed chromatin is strictly dependent on transcription, we monitored FACT/RNApol II ratios by ChIPseq, using anti-SUPT16h and anti-POLR2A antibodies. The results shown in Fig 3A indicate a significant increase of the SUPT16h/POLR2A ratios in the PFDN5-KO metagene that represented the average of all expressed genes, with respect to the wildtype control. This change was not due to a higher overall level of FACT in PFDN5-KO cells, either in chromatin or in the soluble fraction (Fig S2A)

**Figure 3:**
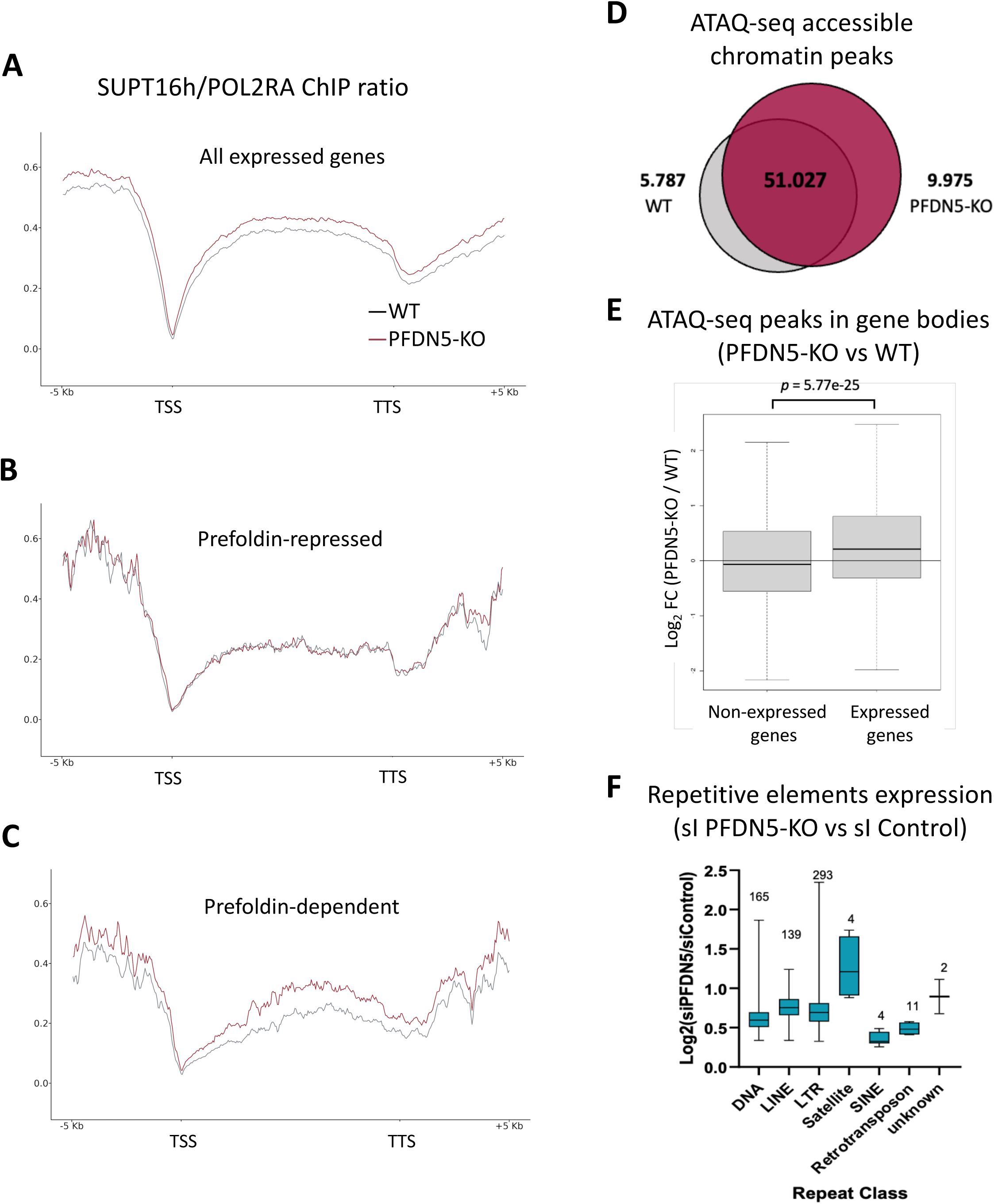
Lack of prefoldin impairs transcriptional dynamics and function of FACT. Metagene analysis on SUPT16h/POL2RA ChiP-seq ratios represented in A) all expressed genes B) prefoldin-repressed genes C) prefoldin-dependent genes, according to how HCT116 genes were affected by prefoldin depletion under serum starvation conditions (Payán-Bravo et al, 2021). Graphs represent the average of two independent paired replicates of SUPT16h and POL2RA ChIPs. D) PFDN5-KO cells show higher frequency of intragenic open chromatin in expressed genes. Number of ATAC-seq peaks detected in wild-type and PFDN5-KO cells. D) Fold-change of ATACseq-detected peaks in PFDN5-KO versus control HCT116 cells, in the bodies of expressed and non-expressed genes. E) Genomic repeat elements get activated upon prefoldin perturbation. Analysis of repetitive elements using the RNA-seq data in si-PFDN5 and si-control cells (Payán-Bravo et al, 2021). Repetitive elements detection was based on subfamily categories. The numbers of repetitive elements used in each category is indicated.

Our previous work had shown that prefoldin provokes different effects across the genome. In some genes, prefoldin depletion produced downregulation in its response to serum starvation, whereas in others the detected output was upregulation [10]. We call prefoldin-dependent to the first subset of genes and prefoldin-repressed to the latter. Our POLR2A ChIP confirmed the opposite transcriptional effect of PFDN5-KO in these two groups of genes, with higher occupancy along the entire metagene in prefoldin-repressed genes, and lower occupancy in the prefoldin-dependent subset (Fig S2B). When we monitored SUPT16H/POLR2A ratios in these two different groups of genes, we found that no alteration was detected in prefoldin-repressed genes, whereas very clear increased values were exhibited by PFDN5-KO cells in the prefoldin-dependent metagene (Fig 3B-C).

As we mentioned in the Introduction, lack of prefoldin does not only cause chromatin-related phenotypes in *Saccharomyces cerevisiae*. We also detected a severe reduction in Ser2-phosphorylation of RNApol II CTD in budding yeast [9] that was later confirmed in human cells [10]. Among the putative interactors of prefoldin in chromatin, we found U2AF2, also called U2AF65 (Fig 2A), an essential factor needed for the co-transcriptional activation of pre-mRNA spliceosomes that specifically binds Ser2-phosphorylated CTD [13]. Since we had already detected defective U2AF65 recruitment to transcribe chromatin under prefoldin perturbation [10], we wondered whether the negative influence of PFDN5-KO on FACT dynamics might be also connected to defective phosphorylation of elongating RNApol II. We calculated the specific levels of phosphorylation by ChIP-seq using an anti-Ser2P antibody in serum-depleted conditions. After normalization with the corresponding anti-POLR2A ChIP-seq signal, we found severely decreased proportion in the all-expressed-genes metagene in PFDN5-KO cells (Fig S2C). However, and in contrast to FACT/RNApolII ratios, this decreased was general and equally found in prefoldin-repressed and prefoldin-dependent genes (Fig S2C), dismissing a relationship with the phenomenon that we address in this work.

A simple interpretation of the specific increase of the FACT/RNApol II ratio in the set of genes that depends on prefoldin for its transcription would be that they require prefoldin for the optimal action of FACT. Accordingly, this dysfunctional FACT of PFDN5-KO cells would stay longer in the vicinity of the transcription site, provoking increased occupancy. Alternatively, prefoldin might be impairing FACT recruitment to transcription elongation sites in this subset of genes. To distinguish between these two options, we investigated whether prefoldin contributed to FACT function in our cells. As described in the Introduction, FACT is not only a factor that facilitates transcription in chromatin, but an integrity factor that ensures nucleosome repositioning after RNApol II passage [11]. To detect possible chromatin integrity defects in the absence of prefoldin, we analysed chromatin accessibility by ATAC-seq. As shown in Fig 3D, we detected around 6x10^4^ peaks in wildtype and PFDN5-KO cells, with very high overlap. Interestingly, when we focused in those accessible peaks located in intragenic regions, we found a very significant difference by PFDN5-KO in expressed *versus* non-expressed genes (Fig 3E), suggesting a chromatin integrity defect linked to transcription elongation.

Defects in chromatin reassembly caused by FACT malfunction has been linked to activation of cryptic transcription initiation sites within the body of actively transcribed genes in budding yeast [14–15]. Moreover, FACT has been shown to repress retrotransposons and cryptic promoters derived from them in mammalian cells [16]. We explored our previous transcriptomic data of prefoldin-depleted cells and found that all the classes of repetitive elements tested, including DNA transposons, LINE, LTR, satellite, SINE and retrotransposons, exhibited increased expression (Fig 3F).

### Lack of prefoldin impairs macroH2A1 transcriptional eviction

The results explained above suggest that prefoldin may stimulate transcription of specific genes by cooperating with FACT in its chromatin functions. As FACT is a chaperone able to interact with all four nucleosomal histones during transcription, we decided to address their dynamics under controlled conditions in an individual model gene by ChIP-qPCR. Me measured histones occupancy in three different positions of *CTNNBL1*, one of the genes whose expression is positively affected by prefoldin (Fig S3A), under chemical control of transcription by DRB. Under transcription inhibition, we did not detect any difference between PFDN5-KO and control cells, for any of the histones tested (Fig 4A). 60 min after washing the drug out, we did not find difference either for the four canonical histones, except H2B, which accumulated significantly in PFDN5-KO cells, in the two more promoter-distal positions tested (Fig 4A). Since H2B is always forming a complex with a H2A-type partner, we explored non-canonical versions of this histone. We found no difference for H2A.Z, which, as expected, was only detected in the most promoter-proximal position. However, we did find significant accumulation of macroH2A1, in PFDN5-KO cells, reproducing the same pattern that H2B (Fig 4A). This result cannot be explained by increased overall levels of any of these histones, since they did not undergo significant differences in PFDN5-KO cells (Fig S3B).

**Figure 4:**
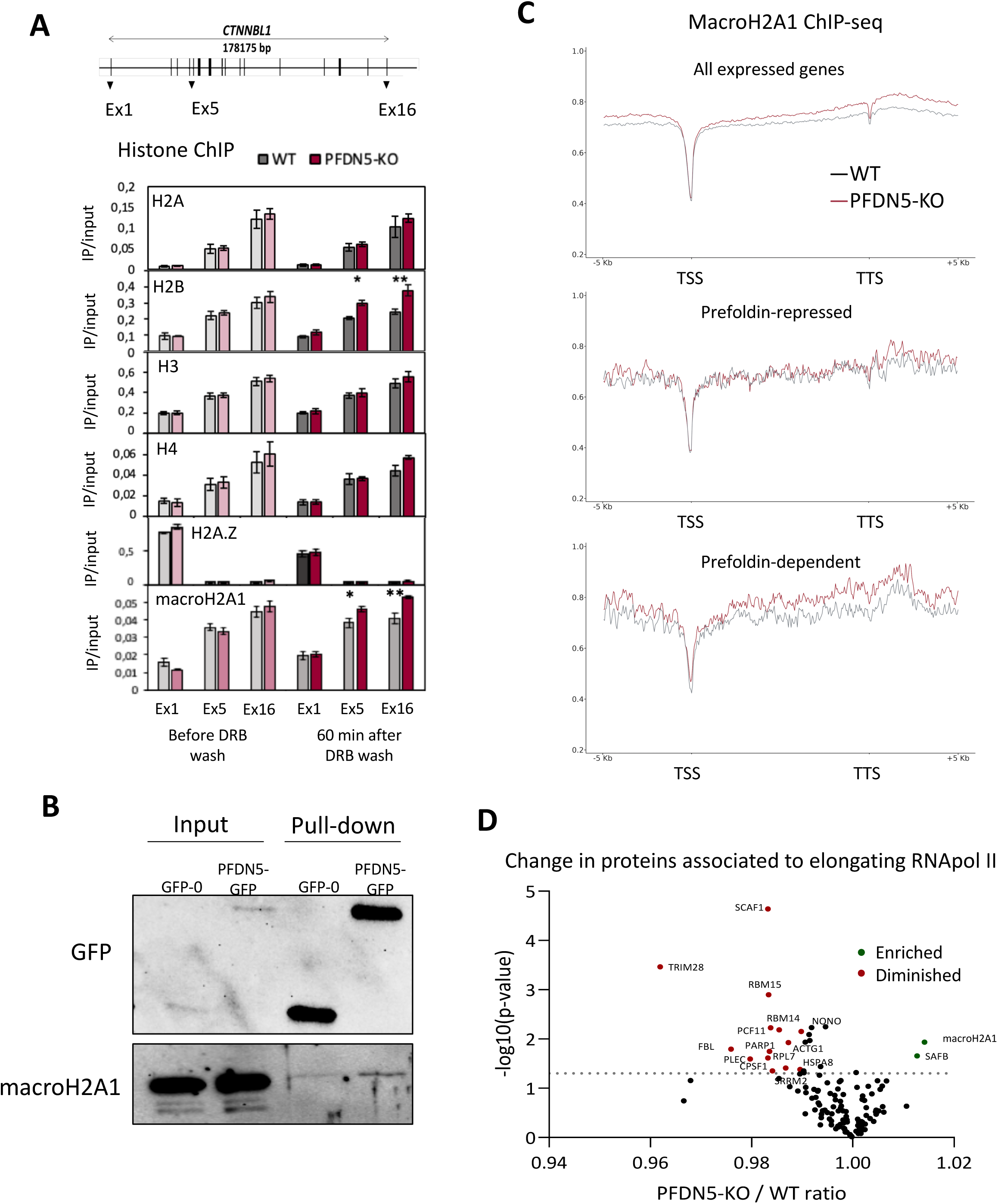
Lack of prefoldin impairs macroH2A1 transcriptional eviction. A) Transcription-dependent accumulation of macroH2A-H2B in the gene body of *CTNNBL1*, in PFDN5-KO cells. ChiP q-PCR analysis of canonical histones H2A, H2B, H3, H4, and non-canonical histone variants H2A.Z and macroH2A1 after 3 hours of 100uM DRB treatment (t=0 min) and 60 minutes after DRB wash. Each immunoprecipitation was normalized against input. The graphs represent the mean and the standard deviation of four independent replicates. * t-test with p-value <0.05, ** t-test with p-value <0.005. B) Physical interaction of prefoldin with macroH2A1 as shown after PFDN5-GFP pull-down experiments. Western blot analysis of macroH2A1 against PFDN5-GFP and GFP pull-down of the chromatin fraction. C) Metagene representation of macroH2A1 ChiP-seq average values in PFDN5-KO (red) or control cells (black), in all expressed, prefoldin-repressed and prefoldin-dependent gene subsets, according to how HCT116 genes were affected by prefoldin depletion under serum starvation conditions (Payán-Bravo et al, 2021). D) MacroH2A1 accumulates at RNApol II elongation complexes in PFDN5-KO cells. Anti-Ser2-phosphorylated CTD IP experiments performed against nuclear extracts of PFDN5-KO and control cells were analysed by iTRAQ mass spectrometry. The proteins detected were filtered by score and normalized by the POL2RA signal of each of the four idependent replicates. Statistical analysis was performed using the Holm-Sidak method. PFDN5-KO/WT protein ratio and -log10 (p-value) were used for representation in a volcano plot. Red dots indicate significantly downregulated proteins, green dots indicate significantly upregulated proteins and black dots indicate non-significant proteins.

MacroH2A1contains a macrodomain connected by an unstructured 41 amino acid long linker that situates it outside of the nucleosomal core [17]. It turned out to be that this H2A-like histone was one of the 35 top interactors of chromatin prefoldin detected in our proteomic screening (Fig 2A). We confirmed this interaction by western blot in new pull-down experiments (Fig 4B). We also performed anti-macroH2A1 ChIP-seq analyses under serum starvation conditions. The results shown in Fig 4C revealed significant effect of PFDN5-KO in the accumulation of macroH2A1 in the metagene of all expressed genes and in the one corresponding to the prefoldin-dependent subset, coinciding with the gene groups that displayed increased FACT/RNA Pol II ratios, but no difference was found in the prefoldin-repressed group of genes (Fig 4C). Quantitation confirmed significant increase of macroH2A1 occupancy in PFDN5-KO cells, in prefoldin-dependent versus prefoldin-repressed genes (Fig S3C). No accumulation was found in the parallel ChIP-seq analyses performed for canonical histones, not even for H2B, likely due to the low proportion of it associated to macroH2A1 in comparison to canonical H2A (Fig S3C-E).

The repressive role described for macroH2A1 in the literature [18] predicts a scenario of RNApol II elongation complexes pausing more often in front of macroH2A1 in PFDN5-KO cells. To address this hypothesis, we performed a second proteomic analysis in which we analysed RNApol II elongation complexes immunoprecipitated from PFDN5-KO solubilized chromatin. The volcano plot in Fig 4D shows the differences found in their composition as compared to control cells. Several CTD-binding proteins involved in cotranscriptional pre-mRNA processing were diminished in the mutant samples, likely as a consequence of the Ser2 phosphorylation defect characteristic of prefoldin-deficient cells (Table S2). On the other side of the plot, only two proteins were significantly increased in PFDN5-KO cells, being macroH2A1 the one exhibiting the highest enrichment (Fig 4D).

The combination of all the results described in this section makes us to conclude that the physical interaction of prefoldin with macroH2A1 in the chromatin context facilitates its eviction during transcription elongation.

### Prefoldin controls FACT and macroH2A1 occupancy in the same subset of genes

We wondered whether the physical and functional interactions detected between prefoldin, on one side, with FACT and macroH2A1, on the other side, were independent or were the consequence of a coordinated action of the three proteins in gene transcription. As a first approach, we measured SUPT16h occupancy alterations in the genes subsets with significant changes of macroH2A1 occupancy. We found that, in average, those genes with increased macroH2A1 levels in PFDN5-KO cells (log_2_FC>2) also exhibited increased SUPT16h occupancy. This change was not observed in genes with decreased macroH2A1 occupancy and was highly significant with respect to the large set of genes with no significant change in macroH2A1 occupancy (Fig 5A). Likewise, the subset of genes with increased SUPT16h occupancy in PFDN5-KO cells (log_2_FC>2) displayed increased levels of macroH2A1, with high significance with respect to the rest of the genome (Fig 5B).

**Figure 5:**
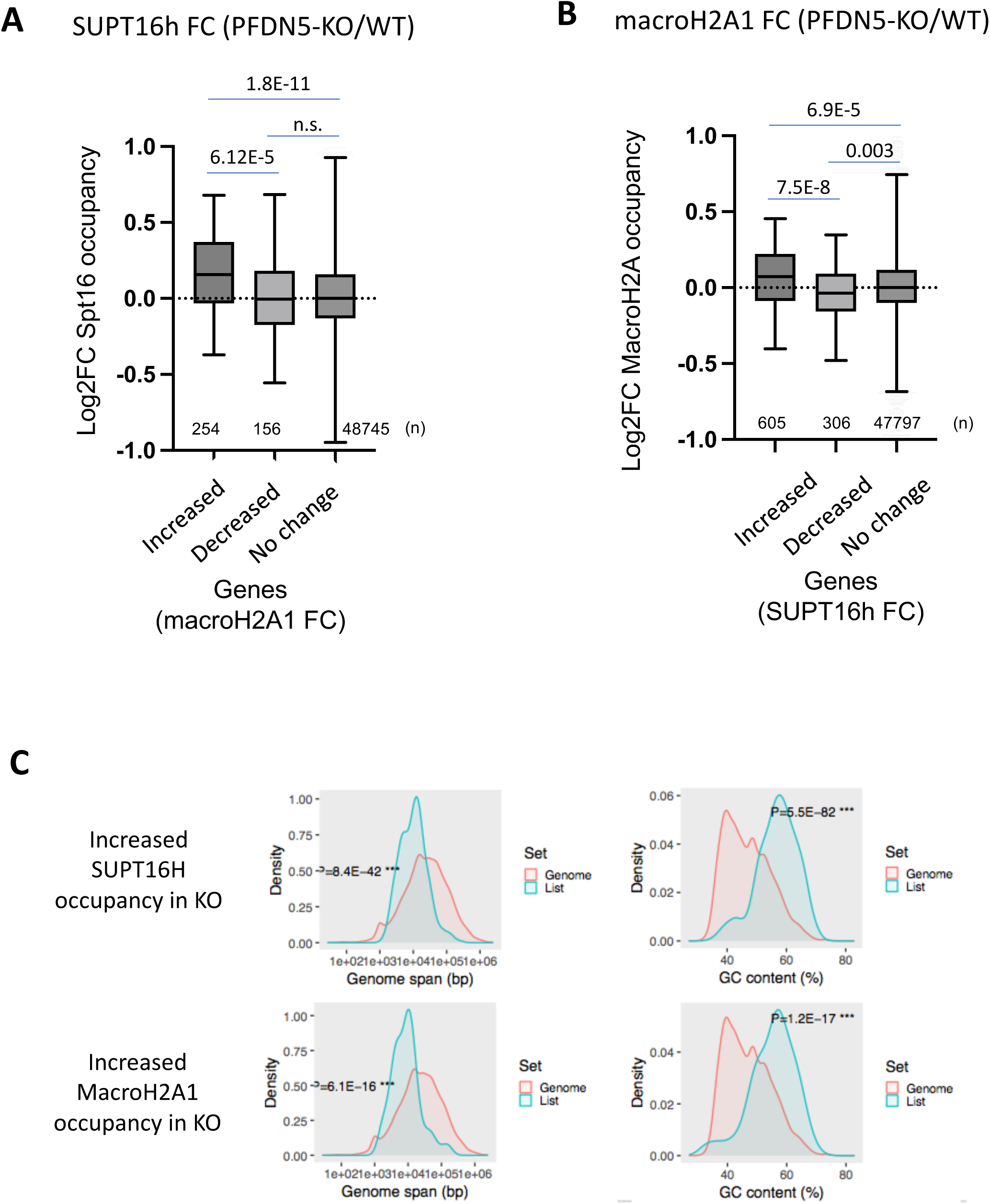
Lack of prefoldin provokes parallel increased occupancy of FACT and macroH2A1 across the genome. A) Change of SUPT16h occupancy in PFDN-KO cells in three subsets of genes, classified according to the effect of PFDN5-KO on the occupancy of macroH2A1 (increased, decrease or no significant change). B) Change of macroH2A1 occupancy in PFDN-KO cells in three subsets of genes, classified according to the effect of PFDN5-KO on the occupancy of SUPT16h (increased, decrease or no significant change). C) Statistical distribution of gene length (Genome span) and GC content in genes with increased occupancies of SUPT16h or macroH2A1, in comparison to the whole genome.

Interestingly, genes with increased SUPT16h or macroH2A1 occupancy in PFDN5-KO cells share common physical features in terms of gene length (shorter than the genome average) and GC content (significantly higher) (Fig 5C). In contrast, genes with lower SUPT16H or macroH2A1 occupancy in PFDN5-KO cells did not exhibit comparably substantial bias (Fig S4).

### Prefoldin is required for FACT-macroH2A1 co-localization in the cell nucleus

The co-regulated occupancy of FACT and macroH2A1 by prefoldin in the same subset of genes suggests that the latter may be facilitating the interaction of the two formers during gene transcription. In order to test this hypothesis, we performed proximity ligation assays (PLA), using specific antibodies against the three proteins. Our results showed some co-localization of prefoldin and macroH2A1 in the nucleus that became significantly enhanced under serum starvation conditions (Fig 6A and S5A). This increase in co-localization after serum starvation was very robust, as indicated by the significant difference between the medians of the three independent replicates that we performed (Fig S5B). We also detected significant co-localization of prefoldin and FACT in the nucleus that was also moderately enhanced by serum starvation, although less robustly (Fig 6B and S5C-D), suggesting that prefoldin-FACT interaction is less influenced by regulatory stimuli.

**Figure 6:**
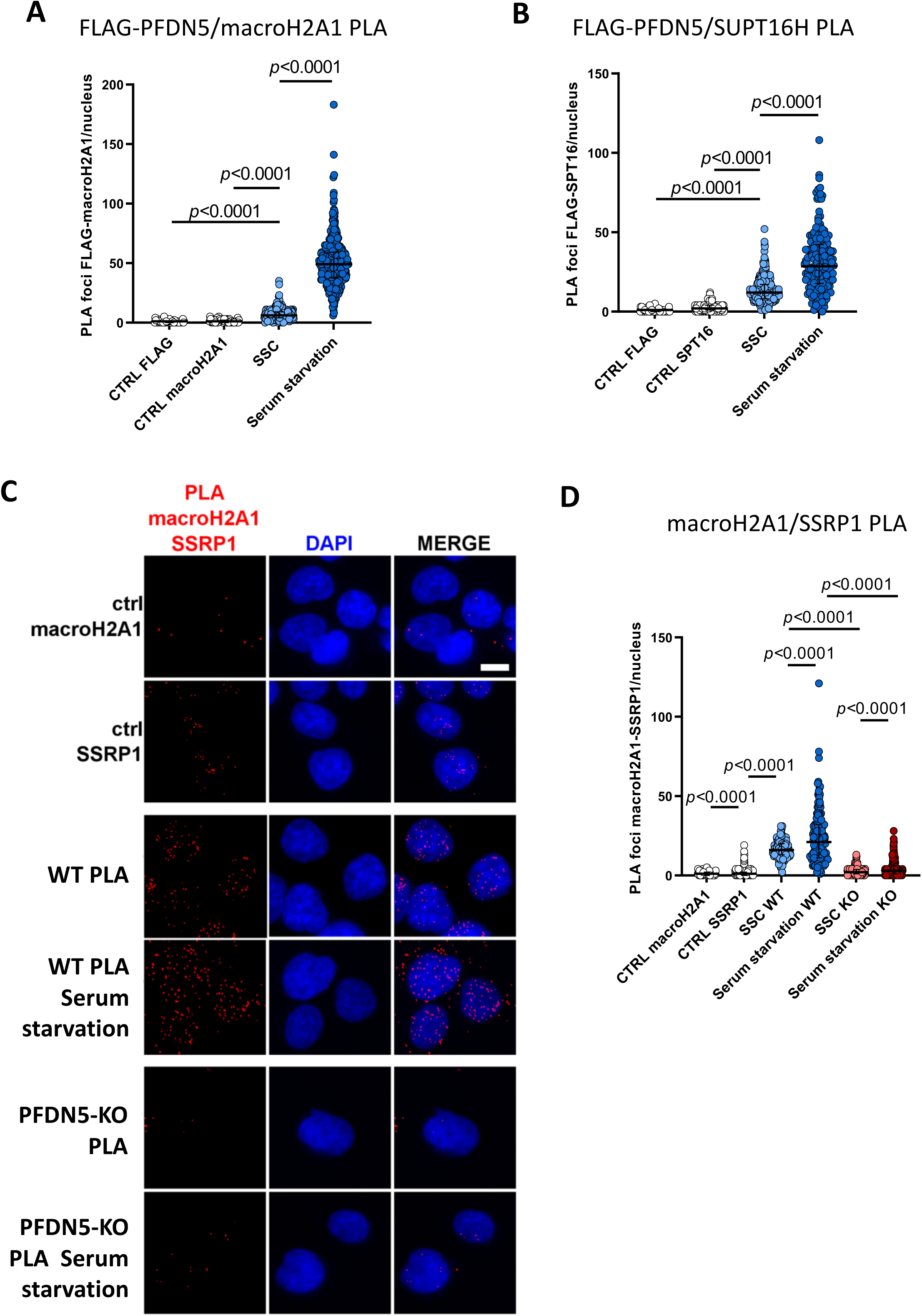
Prefoldin enables FACT-macroH2A1 interaction in response to serum starvation. Proximity-ligation assay results for FLAG-PFDN5 and macroH2A1 (A), FLAG-PFDN5 and SUPT16h (B) or macroH2A1 and SSRP1, in wild-type and PFDN5-KO cells (C, D). Quantification of PLA foci per nucleus of HCT116 cells grown in steady state conditions or after incubation with serum-free culture media for 48 hours. CTRL (PLA with only one primary antibody) are negative controls. Data are the median with interquartile range of all the nuclei analized in three independent experiments (more than 100 nuclei per condition per replicate). Statistical significance was determined by unpaired t-test.

Our PLA results also showed clear proximity between FACT and macroH2A1 in the nuclei (Fig 6C-D and S5E), in agreement with the previously reported role of FACT in macroH2A1 dynamics (Sun et al, 2018; Ji et al, 2024). This FACT-macroH2A1 co-localization, which was also enhanced by serum starvation, was almost abolished in PFDN5-KO cells (Fig 6C-D and S5E).

## Discussion

Prefoldin was described as a co-translational co-chaperone that acts on partially folded nascent polypeptides and delivers them to the ATP-dependent chaperonin TCP for final folding [2]. This cytoplasmic function of prefoldin is particularly essential for proteins with complex folding patterns, such as actin and tubulin monomers [3]. Our previous work has shown that prefoldin is also required for nuclear functions related to gene transcription in both yeast and mammalian cells [9–10]. One of the phenotypes observed in prefoldin mutants is the alteration of histone dynamics during transcriptional elongation [9]. The results of the present study confirm the presence of the canonical prefoldin complex, with its all six subunits (Figs 1B and 2A), within human chromatin and its involvement in transcriptional chromatin dynamics.

Our results further demonstrate that at least one of the mechanisms linking prefoldin to chromatin dynamics is its ability to physically interact with FACT. Prefoldin and FACT mutants share phenotypes characterized by compromised chromatin integrity and the activation of cryptic transcription (Fig 3D-F) [11, 14–15]. In cells lacking prefoldin, the local presence of FACT within gene bodies is altered in response to serum deprivation; this occurs specifically within the subset of the genome whose upregulation by this stimulus is positively dependent on prefoldin (Fig 3A-C) [10]. We therefore conclude that prefoldin’s contribution to the transcriptional induction of this set of genes lies, at least in part, in its physical and functional interaction with FACT.

FACT is the primary histone chaperone that operates during transcription elongation [11]. One of its functions is the selective eviction of macroH2A1 [19], a non-canonical H2A-like histone variant that exerts a repressive effect on transcription [18]. Our results show that prefoldin physically interact with macroH2A1, and its absence alters the co-transcriptional dynamics of this histone (Fig 4-B). Furthermore, lack of prefoldin leads to selective accumulation of this histone in response to serum deprivation, specifically at the same set of genes whose transcription depends on prefoldin, and where FACT also accumulates (Fig 4C). Moreover, the accumulation of macroH2A1 at a gene in response to the absence of prefoldin predicts the accumulation of FACT, and vice versa (Fig 5A-B), which suggests that all three proteins participate in the same molecular process required for the proper transcriptional regulation of these genes.

MacroH2A1 expresses in two different splice variants: macroH2A1.1 contains a functional ligand-binding pocket in its macrodomain capable of binding ADP-ribose and related NAD+-derived metabolites that is lacking in macroH2A1.2 [12]. This second variant is the most abundant in the cell line used for this work. It has been described that macroH2A1.1 contributes to the transcriptional regulation of target genes in two distinct ways. It can do so either through its association with the transcription start site (TSS) of genes characterized by paused RNA Pol II, or via a more general repressive function, shared with macroH2A1.2, that acts upon transcriptional elongation across the gene body [20]. Our results suggest that prefoldin cooperates with FACT in regulating the dynamics of this histone in the context of this second repressive function, since in the absence of prefoldin it does not accumulate at the TSS, but rather spreads across the entire length of the gene (Fig. 4A,C).

Our previous work had shown that the absence of prefoldin leads to severe defects in the Ser2 phosphorylation of RNA Pol II [9–10] and that this results in drastic alterations of pre-mRNA co-transcriptional splicing. This, in turn, affects genome expression in a non-specific manner, with a particularly pronounced effect on long genes containing a high number of introns [10]. However, the results of the present study demonstrate that the impact of prefoldin deficiency on FACT-mediated macroH2A1 dynamics occurs specifically in genes that require prefoldin for their regulation under serum-deprivation conditions. Notably, these genes tend to be shorter than the genomic average (Fig. 5C). Furthermore, the Ser2 phosphorylation defect resulting from the absence of prefoldin is not limited to genes that are positively regulated by prefoldin in response to serum deprivation but very general (Fig. S2C). We therefore conclude that the role of prefoldin in FACT-mediated chromatin dynamics is not a consequence of its function in RNA Pol II phosphorylation.

Prefoldin contribution to chromatin dynamics may be directly related to its chaperone function [21], though not necessarily in cooperation with the CCT/TRiC complex, as occurs when it acts co-translationally in the cytoplasm [1–3]. The CCT/TRiC has been recently described to be also present in the cell nucleus [22], in association to prefoldin [23]. However, none of the putative targets of nuclear CCT/TRiC coincides with FACT or macroH2A1, or is contained in the list of putative prefoldin interactors identified in this work [23]. In fact, we did not detect TCP-1 association with prefoldin in our chromatin preparations (Figs 1B, 2A).

If prefoldin is acting as a chaperone in the FACT-macroH2A1 scenario, a pending question is which of the two proteins would be its chaperone client. The repressive effects of macroH2A1 stem primarily from the properties of its non-histone region, which localizes outwards of the nucleosomal core, stabilizes DNA at its entry/exit site and promotes self-association of nucleosomal arrays [24, 25]. Furthermore, it has been shown that this region blocks the ability of FACT to maintain the integrity of macroH2A1-containing nucleosomes during transcriptional elongation, leading to their eviction [26]. Given that prefoldin appears to be specifically required for FACT to interact efficiently with macroH2A1 (Fig 6C–D) and that both FACT and macroH2A1 co-localize with prefoldin (Fig 6A-B), we propose that the latter would act as a bridging element, connecting FACT and macroH2A1. In this context, prefoldin action on the non-histone region of macroH2A1 would modify its folding in a manner that would allow FACT to engage with the nucleosomal core. Prefoldin might also modulate FACT mode of action, shifting it from chromatin integrity into histone eviction. Future work will be needed to elucidate these hypotheses.

## Materials and methods

### Cell culture

The human colorectal carcinoma cell line HCT116 was cultured in McCoy’s 5A modified medium supplemented with 10% (v/v) fetal bovine Serum (FBS), 60 mg*/*l penicillin, and 100 mg*/*l streptomycin at 37°C and 5% of CO_2_. Additionally, 2 g/l G418 were added to resistant cell lines HCT116 PFDN5-Flag, HCT116 Flag, HCT116 GFP-0, HCT116 PFDN5-GFP, HCT116 pCDNA3.1 GFP and HCT116 pcCDNA3.1 SSRP1-GFP.

### DNA transfection

HCT116 cell line were seeded one day before transfection at a density of 1 x10^5^ cells in each well of a 6-well dish. 6 µl of Turbofec^®^, 2 µg of plasmid of interest and 100 µl of optiMEM were mixed using vortex. After the incubation of 15 min at RT, the solution was added dropwise to the cells. One day after transfection, 2g/l G418 were added to the cells. To generate stable cell lines, the cells were maintained with G418, sorted by flow cytometry and checked the construction of interest by western blot.

### Chromatin immunoprecipitation

Approximately 30 x 10^6^ growing cells were cross-linked using formaldehyde to a final concentration of 1% (p/v) in the crosslinking solution (50 mM HEPES pH 8.0, 0.1M NaCl, 1mM EDTA pH 8.0, 0.5mM EGTA pH 8.0) during 10 min at 37°C. In order to stop the cross-linking reaction, 2 M glycine was added to the cells for 5 min at RT. Cells were washed twice with cold PBS 1X and centrifuged 3600 g for 5 min at 4°C. The cell pellet was treated with 2.5 ml of Lysis Buffer 1 (5mM Pipes pH 8.0, 85mM KCl, 0.5% v/v NP-40) with protease inhibitors and incubated for 10 min on ice, after the incubation, lysed cells were centrifuged at 3600 g for 5 min at 4°C, the nuclear cell pellet was resuspended in 1 ml of Lysis Buffer 2 (1% SDS, 10mM EDTA pH 8.0, 50mM Tris-HCl pH 8.0) containing protease inhibitors (1mM PMSF plus Complete protease inhibitor cocktail). Chromatin was then sonicated 30s on and 30s off for 15 min at 4°C to get DNA fragments between 300bp to 500bp. After sonication, chromatin was cleared by centrifugation firstly at 3600 g for 5 min at 4°C and later at 15000 g for 10 min at 4°C.

For each immunoprecipitation, we used 100 µl of chromatin, 900 µl of IP buffer (0.01 % SDS, 1.1% Triton X-100, 1.2mM EDTA, 16.7mM Tris-HCl pH 8.0, 167mM NaCl) containing protease inhibitors and the corresponding antibody (Table S3). The mixture was incubated at 4°C over-night. In the following day, 50µl of magnetic dynabeads Protein A or G was added to each sample and incubated on rotation at 4°C for 2 hours. After the incubation, dynabeads were washed with the wash buffers 1 (1% SDS, 1% Triton X-100, 2mM EDTA, 20mM Tris-HCl pH 8.0, 150mM NaCl), 2 (1% SDS, 1% Triton X-100, 2mM EDTA, 20mM Tris-HCl pH 8.0, 500mM NaCl), 3 (0.25M LiCl, 1% NP-40, 1% NaDoc, 1mM EDTA, 10mM Tris-HCl pH 8.0) and TE buffer (10mM Tris-HCl ph 8.0, 1mM EDTA). Samples were then eluted at 65°C with the elution buffer (10mM Tris-HCl pH 8.0, 1mM EDTA, 1% SDS) for 10 min. For DNA purification, samples were incubated at 65°C over-night, treated with proteinase K (20mg/ml) for 1h at 37°C. The resultant DNA was treated with phenol: chloroform and precipitated with 2.5V of Ethanol, 1/10 (v/v) sodium acetate 3M and 0.1% of glycogen and centrifuged at max speed for 30 min. DNA was finally resuspended in 200µl of miliQ H_2_O.

Real-time quantitative PCR analyses were performed in Light Cycler 480 II thermocycler (Roche). For each reaction we use 384 plates, mixing 4µl of each sample with 6 µl of the mixture 0.5 µl of H_2_O, 0.5 µl Forward and reverse oligonucleotides (10µM) and 5 µl of SYBR Green Premix 2X.

For ChiP-seq experiments, we followed the procedure described above, except that we did not add glycogen for DNA purification. Before immunoprecipitation, 1.5 µg of *S.cerevisiae* chromatin was added to spike-in the samples. DNA purification after proteinase K treatment was performed using the ChIP DNA Clean & Concentrator kit (Zymo Research). All the libraries preparation and sequencing was performed at the Genomic Unit of CABIMER (Seville, Spain), using the ThruPLEX DNA-seq kit and the illumine NovaSeq 6000 instrument.

### Pull down of chromatin fraction using GFP-trap

Cells were seeded around 1x10^6^ cells per 10 cm plate, after 3 days (2 plates per condition) were washed with PBS (138 mM NaCl, 10 mM Na2HPO4, 1.8 mM KH2PO4, 3 mM KCl, pH 7.4), scraped and centrifuged at 900 g for 5 min. The pellet was store at -80°C or lysed with 1ml of IP buffer (50mM Tris-HCl pH 7.5, 150 mM NaCl, 0.5% NP-40, 2mM MgCl_2_) containing protease inhibitors, and maintained for 20 min on ice. To get the chromatin fraction, cells were centrifuged at maximum speed for 10 min at 4°C. The supernatant was removed and the pellet was resuspended in 1ml of the same buffer and incubated with 750 U of Benzonase for 3 hours at 4°C in rotation.

To clarify the lysate, the sample was centrifuge at 110,000 g for 60 min at 4°C, then 100 µl of this extraction was taken as input, 25 µl of equilibrated agarose GFP-trap beads were added for the rest of the sample that was incubated for 90 min at 4°C on rotation. Finally, beads were washed six times with the IP buffer and the chromatin protein bound to the beads was eluted with 100 µl of Laemli 2x solution (Tris-HCl pH 6.8 160 mM; glicerol 20 %; SDS 4 %; DTT 0.2M and bromophenol blue 1 %) at 100°C for 5 min.

For mass spectrometry analysis, samples were eluted in 50µl of acidic buffer (200Mm Glycine pH 2.5) pipetting for 30s, samples were later neutralised with 5ul of 1M Tris-HCl pH 10.4. MS/MS tandem mass spectrometry though Label-Free Quantitation (LFQ) was performed.

### iTRAQ

The iTRAQ experiment was performed as followed. Four independent replicas of wild type and PFDN5-KO cells were grown and collected under steady state condition. All replicas were extracted with the Nuclear Co-IP kit and immunoprecipitated using RNA Pol II Ser2-P antibody. The same amount of eluted proteins was trypsinised and labelled with isobaric iTRAQ-8plex reagents. After labelling, peptides from all replicas were mixed in the same tube to be analysed by tandem mass-spectrometry analysis.

### ATAC-seq

60,000 cells were harvested per sample and centrifuged at 500 g for 5 minutes at 4°C. The supernatant was carefully removed, the pellet was washed with 200µl of ATAC RSB resuspension buffer (10mM Tris-HCl pH 7.4, 10mM NaCl and 3mM MgCl_2_), and cells were centrifuged at 500 g for 5 minutes at 4°C. The pellet was resuspended gently pipetting up and down three times with 50µl of ATAC RSB resuspension buffer containing 0.1% NP-40, 0.1% Tween 20 and 0.01%. The mixture was then incubated on ice for one minute, followed by an immediate addition of 200 µl of ATAC RSB resuspension buffer containing 0.1% Tween 20. With this procedure cells were lysed to obtain the nuclei that were collected after 10 min of centrifugation at 500G at 4°C. For nuclei tagmentation, nuclei pellets were resuspended in 2.5 µl of 4X Tagmentation Buffer (2X Cut Smart Buffer, 35.2 % PEG 8000), 5 µl of homemade loaded Tn5 and 2.5 µl of H_2_O, and incubated at 37°C for 30 minutes.

Before tagmentation, transposase Tn5 had to be loaded with specific primers. at a final concentration of 45µM. For Tn5 loading, Tn5 transposase and annealed primers were mixed at a concentration ratio of 4:1 and incubated for 1h at 25°C.

For library preparation, tagmentated DNA were subjected to PCR using 15µl of DNA, 2.5 µl of each nextera primer, compatible with the annealed primers of Tn5 at 25µM, 5µl of H_2_O and 25 µl of NEBNext master mix 2x. DNA was later cleaned using AMPure 1.1x and sequenced.

### PLA

PLA experiments were performed as in [27], using the Duolink in situ ddetection kit. Pairs of affinity probes—consisting of specific antibodies coupled to oligonucleotides (Table S3)—were used to detect the colocalization of prefoldin with FACT, prefoldin with macro-H2A1, and FACT with macro-H2A1. An interaction signal is generated only when the proteins are within 60 Å of each other, which enables the hybridization of a connector, enzymatic ligation, and subsequent PCR amplification of a reporter DNA sequence.

### Computational methods and statistical analysis

Most analyses were performed using R version 4.3.1 and RStudio version 1.3.1093.

ATAC-seq data were processed (trimmed, aligned, filtered, and quality controlled) for each condition independently, using the Kundaje laboratory processing package (https://github.com/ENCODE-DCC/atac-seq-pipeline), with the reference human genome hg38. Peak calls were filtered based on the Irreproducible Discovery Rate (IDR) between the two independent biological replicates within each condition. Peaks from both conditions were then merged to generate a consensus peak set. Reads overlapping the hg38 blacklist regions and non-standard chromosomes were removed from downstream analysis. Read counts across consensus peaks were obtained using the regionCounts() function from the *csaw* package (v 1.34.0) (Lun & Smyth, 2015) with minq = 40 parameter. Low-abundance peaks (average logCPM < 0.2) were excluded. For normalization, genome-wide background reads were quantified using windowCounts() with bin = TRUE, width = 2000, and minq = 40 options. Normalization factors were calculated using the TMM method with loess correction implemented in *csaw*. Differential accessibility between the wild-type and knockout conditions was assessed using the Bioconductor package *edgeR* (v 3.42.4) (Robinson et al, 2010), fitting a negative binomial generalized linear model and performing likelihood ratio tests.

ChIP-seq reads from WT and PFDN5-KO samples for Spt16, Rpb1, Ser2P, MacroH2A1, H2A, H2B, and H3 were aligned to the human hg38 reference genome using the function align() from the Bioconductor package *Rsubread* 214 (v2.14.2), with the parameters type=1, TH1=2, and unique=TRUE. For spike-in normalization, reads were also aligned to the Saccharomyces cerevisiae sacCer2 reference genome. The resulting BAM files were sorted and indexed using the Bioconductor package *Rsamtools* (v2.16.0).

ChIP-seq signal profiles were analysed across gene sets comprising upregulated genes, downregulated genes, and all expressed genes in HCT116 cells, as defined in Payán-Bravo et al (2021). Signal matrices were generated using depTools computeMatrix in scale-regions mode, with –m 10000 –bs 50 –a 5000 –b 5000 options. Mitochondrial genes were excluded from the analysis.

Mean signal profiles were subsequently calculated for each gene set. The resulting signal profiles were normalized by multiplying the signal values by the corresponding spike-in normalization factors, obtained from the mapping statistics. Density plots were constructed with *ggplot2* (v3.4.3).

To quantify transposable element (TE) expression, RNA-seq data from Payán-Bravo et al (2021) for siPFDN5 and siControl samples were re-aligned to the human reference genome (hg38) using *STAR*, allowing multimapping reads (up to 100 genomic locations per read). Parameters were as follows: --winAnchorMultimapNmax 100 -- outFilterMultimapNmax 100 --outFilterMismatchNmax 999 --outMultimapperOrder Random --runRNGseed 777--outSAMmultNmax 1 --outFilterType BySJout -- alignSJoverhangMin 8 --alignSJDBoverhangMin 1 --alignIntronMin 20 -- alignIntronMax 1000000 --alignMatesGapMax 1000000.

TE expression was quantified using *TEtranscripts* in multi-mapping mode (--mode multi), comparing the siPFDN5 against siControl conditions, with two biological replicates per condition. Gene and TE annotations were obtained from GENCODE v44 and the RepeatMasker-derived hg38 TE annotation (hg38_rmsk_TE.gtf), respectively.

Differential TE expression analysis was performed using the Bioconductor package *DESeq2*. Count matrices generated by *TEtranscripts* were filtered to retain only features with at least two reads in at least one sample. Differential expression was assessed using a negative binomial generalized linear model. TEs with an adjusted *p*-value < 0.05 were considered for representation.

## Acknowledgments

Conceptualization, S.C., J.R., M.M-C.; Investigation, S.F., E.S-E., E.G., R.C. and A.G-M.; Writing—Original Draft, S.C.; Writing—Review & Editing, S.F., J.R. and M.M-C; Funding acquisition, S.C. and M.M-C.; Supervision, X.P., V.P., S.C., J.R. and M.M-C.

We thank all the people of the IBiS Gene Expression lab for helpful discussion and all technicians from IBiS, CABIMER and CITIUS facilities. We also thank our colleagues from the Gene Expression Lab at IBiS and the Genetics Department of Universidad de Sevilla for their support.

This publication is part of the project PID2023-148037NB-C21, funded by MICIU/AEI/10.13039/501100011033 and by ERDF/EU, to S.C. and M.M-C.

## Data Availability

Sequencing data have been uploaded to the Gene Expression Omnibus (GEO) in a private submission.

## Ethical considerations

This work did not require ethical approval from a human subject or animal welfare committee.

## Supplementary materials

**Figure S1:**
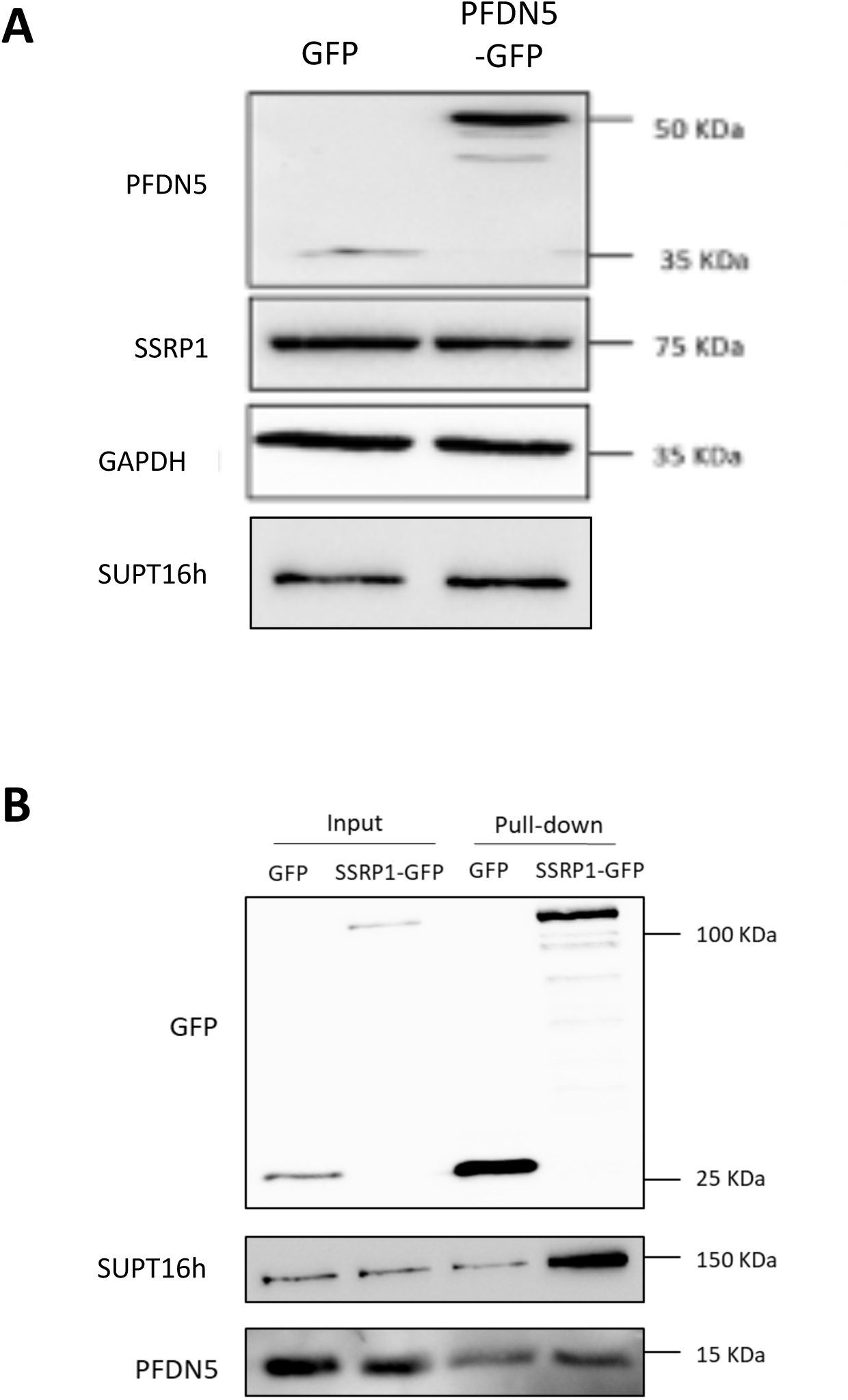
PFDN5 is enriched after SSRP1-GFP pull-down. A) The total protein level of FACT subunits is not influenced by GFP. Western blot experiment of SSRP1 in GFP and PFDN5-GFP human cell constructs. B) Western blot analysis of SSRP1-GFP and GFP pull-down of the chromatin fraction. The membranes were incubated with GFP antibody as a purification control, which showed bands at around 30 KDa for the GFP protein and an upper band at around 120 KDa for SSRP1-GFP. In addition, SUPT16h protein was chosen as a positive control.

**Figure S2:**
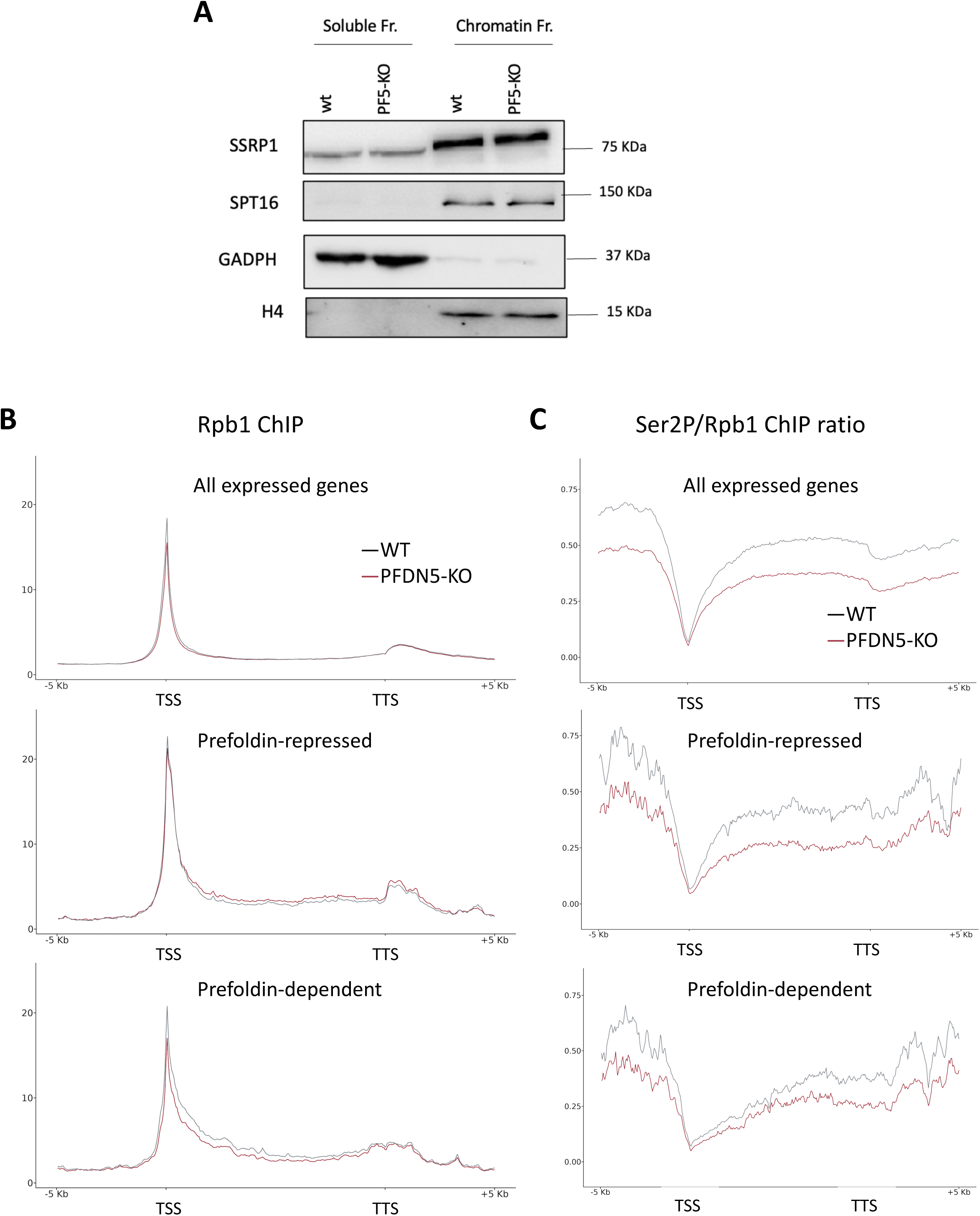
Influence of prefoldin on FACT and macroH2A1 dynamics across the genome does not follow alterations of RNApol II Ser2-phosphorylation. A) The total protein level of the FACT complex is not affected by prefoldin perturbation. Western blots of SSRP1 and SUPT16h in soluble and chromatin fractions of wild-type and PFDN5-KO cells. B) Metagene representation of POL2RA ChiP-seq ratios represented in all expressed, prefoldin-repressed and prefoldin-dependent gene subsets, according to how HCT116 genes were affected by prefoldin depletion under serum starvation conditions (Payán-Bravo et al, 2021). C) Lack of prefoldin impairs Ser2 phosphorylation of RNApol II across the genome, independently of the influence of prefoldin on gene expression under serum starvation. Metagenes represent Ser2-phosphorylated CTD/POL2RA average ChiP-seq ratios calculated from two independent paired replicates of individual ChIPs.

**Figures S3:**
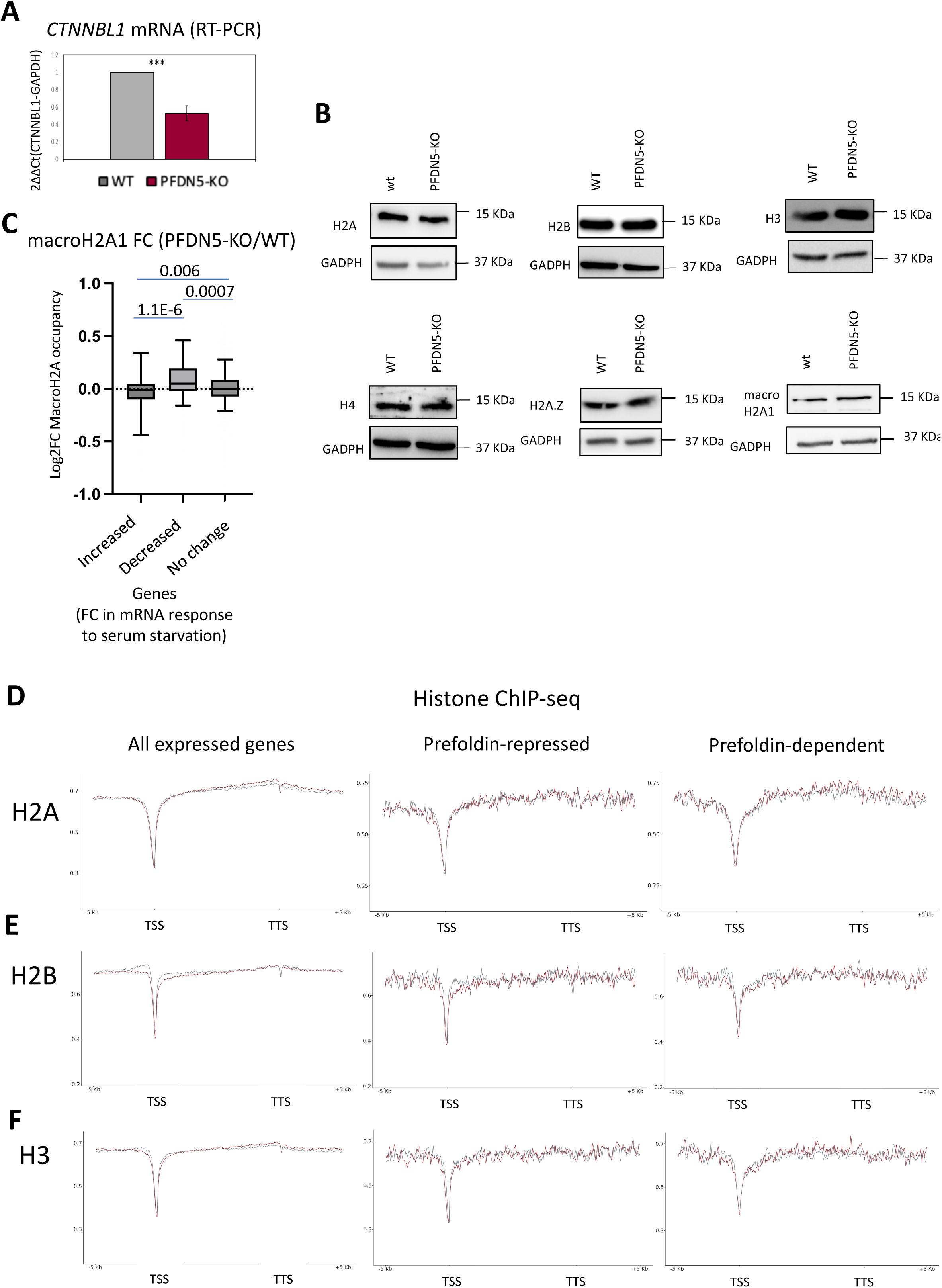
Canonical histones do not accumulate in the chromatin of PFDN5-KO cells in response to serum starvation. A) RT-PCR analysis of mRNA shows that *CTNNBL1* is a prefoldin-dependent gene under standard culture conditions. B) Lack of prefoldin does not alter the total levels of histones. Western blots of total cell extracts, from wild-type and PFDN5 KO cells, performed with antibodies against H2A, H2B, H3, H4, H2A.Z and macroH2A1, and against GAPDH as a loading control. C) Change of macroH2A1 occupancy in PFDN-KO cells in three subsets of genes, classified according to the effect of PFDN5-KO on its mRNA response to serum starvation (increased, decrease or no significant change). D, E and F) Metagene representations of H2A (D), H2B (E) and H3 (F) ChiP-seqs average values in PFDN5-KO (red) or control cells (black), in all expressed, prefoldin-repressed and prefoldin-dependent gene subsets, according to how HCT116 genes were affected by prefoldin depletion under serum starvation conditions (Payán-Bravo et al, 2021).

**Figures S4:**
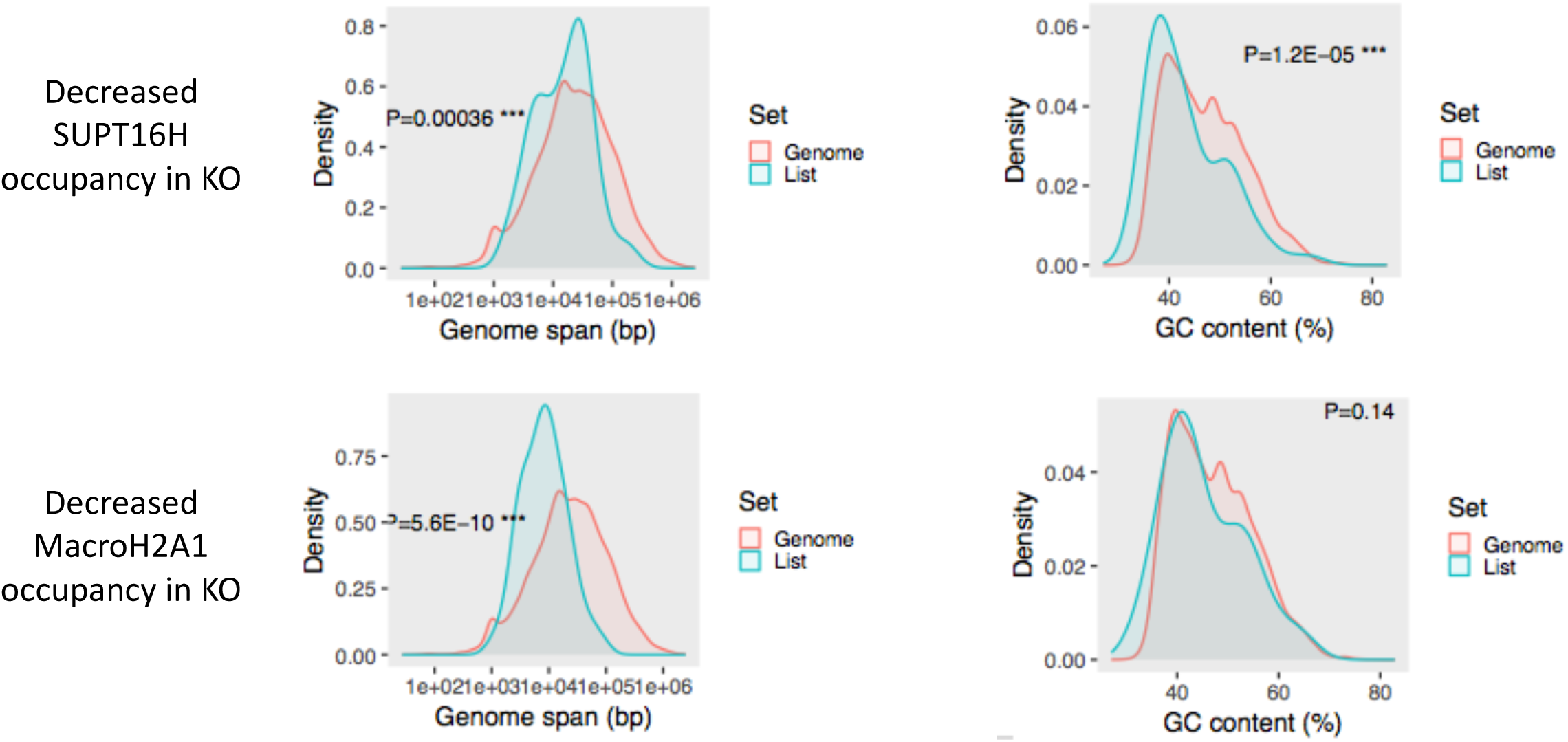
Statistical distribution of gene length (Genome span) and GC content in genes with decreased occupancies of SUPT16h or macroH2A1, in comparison to the whole genome.

**Figures S5:**
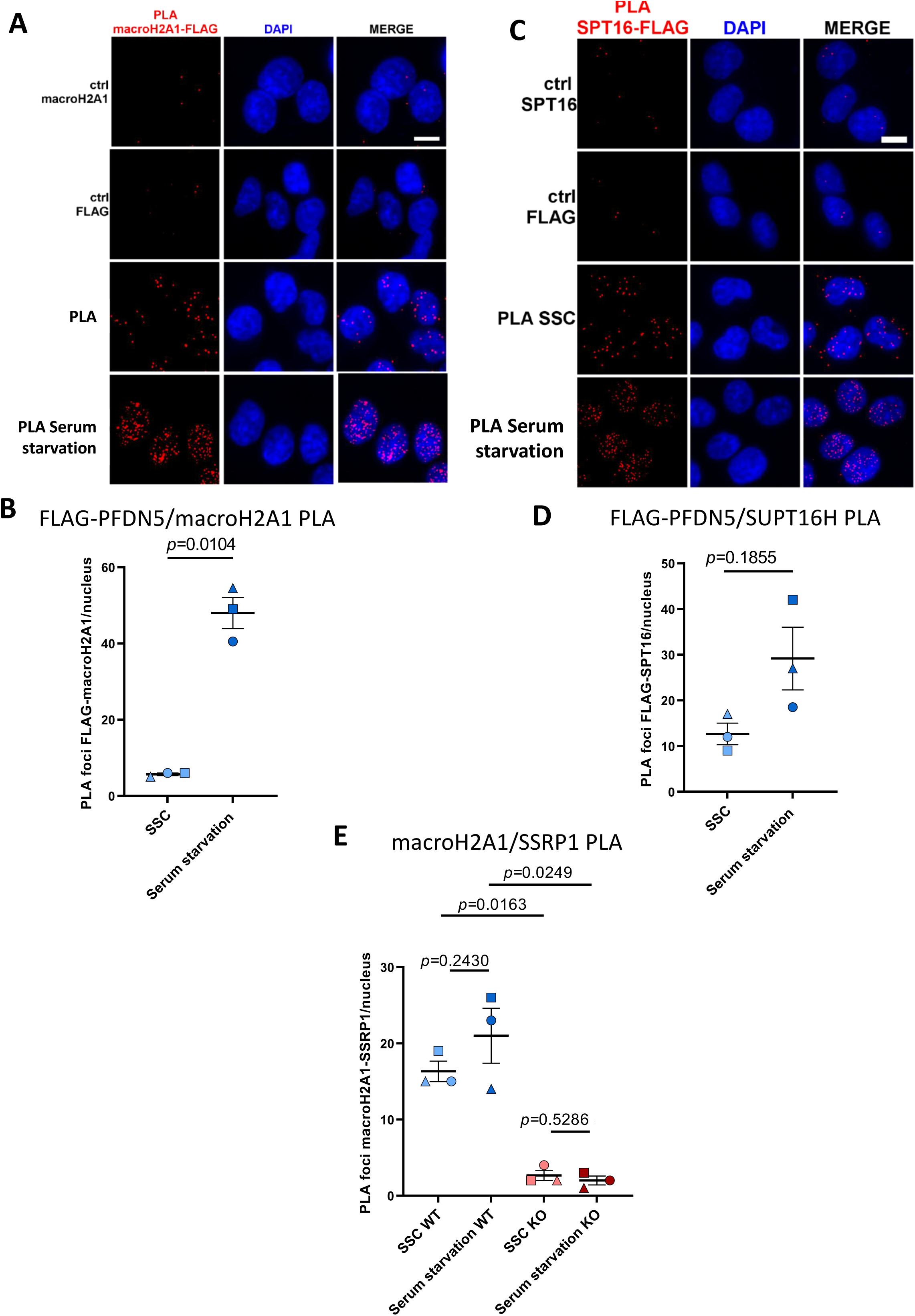
Prefoldin enables FACT-macroH2A1 interaction in response to serum starvation. Proximity-ligation assay results for FLAG-PFDN5 and macroH2A1 (A and B), FLAG-PFDN5 and SUPT16h (C and D) or macroH2A1 and SSRP1, in wild-type and PFDN5-KO cells (E). Quantification of PLA foci per nucleus of HCT116 cells grown in steady state conditions or after incubation with serum-free culture media for 48 hours. CTRL (PLA with only one primary antibody) are negative controls. Data are the mean ± SEM (n=3). Statistical significance was determined by paired t-test.

## Supplementary Tables

**Table S1:** Gene ontology analysis of PFDN5-GFP pull-down results.

| Molecular function categories | Enrichment p-value |
|---|---|
| RNA binding | 1.76E-66 |
| mRNA binding | 5.54E-14 |
| Catalytic activity, acting on a nucleic acid | 2.02E-13 |
| Catalytic activity, acting on RNA | 2.93E-10 |
| Helicase activity | 3.10E-10 |
| Nucleosome binding | 4.02E-08 |
| Nucleosomal DNA binding | 4.02E-08 |
| Cadherin binding | 4.02E-08 |
| Chromatin DNA binding | 1.80E-07 |
| Single-stranded RNA binding | 1.88E-07 |
| DNA replication origin binding | 2.09E-07 |
| Unfolded protein binding | 2.46E-07 |

**Table S2:** Proteins associated to RNA Pol II elongation complexes that become significantly altered in PFDN5-KO cells.

| Protein names | Gene names | Ratio PFDN5-KO / WT | *p* value | Mean Intensity |
|---|---|---|---|---|
| Core histone macro-H2A.1 | H2AFY | 1.014173739 | 0.011599 | 15.46 |
| Scaffold attachment factor B1 | SAFB | 1.012705055 | 0.022041 | 15.54 |
| Non-POU domain-containing octamer-binding protein | NONO | 0.989806377 | 0.007026 | 14.77 |
| Heat shock cognate 71 kDa protein | HSPA8 | 0.989580082 | 0.041476 | 16.72 |
| Actin | ACTG1 | 0.987267551 | 0.011783 | 17.26 |
| Serine/arginine repetitive matrix protein 2 | SRRM2 | 0.986739263 | 0.038933 | 12.21 |
| RNA-binding protein 14 | RBM14 | 0.985460872 | 0.006502 | 14.83 |
| Cleavage and polyadenylation specificity factor subunit 1 | CPSF1 | 0.984090654 | 0.044569 | 13.80 |
| Pre-mRNA cleavage complex 2 protein Pcf11 | PCF11 | 0.983760215 | 0.005904 | 16.65 |
| Poly [ADP-ribose] polymerase 1 | PARP1 | 0.983538304 | 0.017895 | 14.63 |
| Putative RNA-binding protein 15 | RBM15 | 0.983359679 | 0.00127 | 16.76 |
| 60S ribosomal protein L7 | RPL7 | 0.983232599 | 0.024274 | 14.97 |
| Splicing factor, arginine/serine-rich 19 | SCAF1 | 0.983229575 | 0.000023 | 18.78 |
| Plectin | PLEC | 0.979707647 | 0.025525 | 12.81 |
| rRNA 2-O-methyltransferase fibrillarin | FBL | 0.975948339 | 0.01608 | 15.31 |
| Transcription intermediary factor 1-beta | TRIM28 | 0.961921273 | 0.000342 | 12.76 |

**Table S3:** Antibodies used in this work.

| ANTIBODIES | ORIGEN | REFERENCE | ChIP | WB | PLA |
|---|---|---|---|---|---|
| GADPH | Mouse monoclonal | Santa Cruz 47724 | - | 1:1000 | - |
| PFDN1 | Rabbit monoclonal | Abcam 151708 | - | 1:500 | - |
| PFDN2 | Rabbit monoclonal | Bethyl A304-807A | - | 1:100 | - |
| PFDN3 | Mouse polyclonal | Abcam 96085 | - | 1:500 | - |
| PFDN4 | Mouse polyclonal | Abcam 168438 | - | 1:200 | - |
| PFDN5 | Mouse monoclonal | Santa Cruz 271150 | - | 1:100 | - |
| PFDN6 | Mouse monoclonal | Santa Cruz 376733 | - | 1:100 | - |
| FLAG | Mouse monoclonal | Sigma F1804 | - | - | 1:500 |
| GFP | Mouse Monoclonal | Roche 11814460001 | - | 1:5000 | - |
| Tubulin | Rabbit polyclonal | Abcam Ab15568 | - | 1:5000 | - |
| RNA Pol II CTD repeat (phospho S2) | Rabbit polyclonal | Abcam Ab5095 | 5µg | 1:1000 | - |
| 8WG16 | Mouse monoclonal | Thermo MA1-10882 | 4µg | 1:500 | - |
| Rpb1 (D8L4Y) | Rabbit monoclonal | Cell Signalling 14958S | 1µg | 1:1000 | - |
| H2A | Rabbit polyclonal | Abcam Ab18255 | 2µg | 1:1000 | - |
| H2B | Rabbit polyclonal | Abcam ab1790 | 1µg | 1:1000 | - |
| H4 | Rabbit polyclonal | Abcam Ab10158 | 5µg | 1:1000 | - |
| H3 | Rabbit polyclonal | Abcam Ab1791 | 3µg | 1:1000 | - |
| H2A.Z | Rabbit polyclonal | Abcam ab4174 | 3 µg | 1:1000 | - |
| macroH2A1 | Rabbit polyclonal | Abcam ab37264 |  | 1:1000 | 1:500 |
| SUPT16h (12181S) | Rabbit monoclonal | Cell Signaling 12191 | 5µg | 1:1000 | 1:500 |
| SSRP1 | Mouse monoclonal | Biolegend 609702 | 2µg | 1:1000 | 1:500<br>- |
| TCP-1 (H1) | Mouse monoclonal | Santa Cruz sc-137092 | - | 1:500 | - |
| Peroxidase AffiniPure Goat Anti-Mouse IgG | Goat | Jackson 115-035-146 | - | 1:5000 | - |
| Peroxidase AffiniPure Goat Anti-Rabbit IgG | Goat | Jackson 111-035-144 | - | 1:5000 | - |

